# Functional Magnetic Resonance Spectroscopy of Prolonged Motor Activation using Conventional and Spectral GLM Analyses

**DOI:** 10.1101/2024.05.15.594270

**Authors:** Maria Morelli, Katarzyna Dudzikowska, Dinesh K. Deelchand, Andrew J. Quinn, Paul G. Mullins, Matthew A. J. Apps, Martin Wilson

## Abstract

**Background:** Functional MRS (fMRS) is a technique used to measure metabolic changes in response to increased neuronal activity, providing unique insights into neurotransmitter dynamics and neuroenergetics. In this study we investigate the response of lactate and glutamate levels in the motor cortex during a sustained motor task using conventional spectral fitting and explore the use of a novel analysis approach based on the application of linear modelling directly to the spectro-temporal fMRS data.

**Methods:** fMRS data were acquired at a field strength of 3 Tesla from 23 healthy participants using a short echo-time (28ms) semi-LASER sequence. The functional task involved rhythmic hand clenching over a duration of 8 minutes and standard MRS preprocessing steps, including frequency and phase alignment, were employed. Both conventional spectral fitting and direct linear modelling were applied, and results from participant-averaged spectra and metabolite-averaged individual analyses were compared.

**Results:** We observed a 20% increase in lactate in response to the motor task, consistent with findings at higher magnetic field strengths. However, statistical testing showed some variability between the two averaging schemes and fitting algorithms. While lactate changes were supported by the direct spectral modelling approach, smaller increases in glutamate (2%) were inconsistent. Exploratory spectral modelling identified a 4% decrease in aspartate, aligning with conventional fitting and observations from prolonged visual stimulation.

**Conclusion:** We demonstrate that lactate dynamics in response to a prolonged motor task are observed using short-echo time semi-LASER at 3 Tesla, and that direct linear modelling of fMRS data is a useful complement to conventional analysis. Future work includes mitigating spectral confounds, such as scalp lipid contamination and lineshape drift, and further validation of our novel direct linear modelling approach through experimental and simulated datasets.

## 1. Introduction

Functional Magnetic Resonance Spectroscopy (fMRS) is an increasingly popular technique to measure dynamic changes in metabolite levels in response to stimuli (Mullins, 2018; Stanley & Raz, 2018). Whilst the method suffers from poorer sensitivity, and therefore spatial resolution compared to functional MRI (fMRI), it offers a uniquely direct insight to the metabolic processes implicated in neuronal activation. One of the earliest ^1^H fMRS studies demonstrated a significant elevation in visual cortex lactate levels following prolonged (12 minutes and longer) visual stimulation at a field strength of 2.1 Tesla (Prichard et al., 1991). These findings have since been replicated by other groups, using various MRS acquisition techniques and field strengths (Bednařík et al., 2015; Maddock et al., 2006; Mangia, Tkác, Gruetter, et al., 2007; Mangia, Tkác, Logothetis, et al., 2007; Sappey-Marinier et al., 1992; Schaller et al., 2013), with reported participant average increases in lactate levels varying between 19% and 100% of resting levels.

The role of lactate in the brain is directly linked to the metabolism of circulatory glucose to meet neuronal energy demands. Whilst there is general agreement on glucose acting as the primary energy substrate for brain tissue in the resting state, the energetics of elevated neuronal activity are less well understood. Initially regarded as an unwanted byproduct of anaerobic glycolysis, lactate has been more recently proposed as a significant neuronal energy source and signalling molecule associated with plasticity and excitability (Magistretti & Allaman, 2018). Whilst the true importance of lactate to elevated neuronal activity has been controversial (Díaz-García et al., 2017), recent evidence suggests the primary neuronal energy substrate may change from glucose to lactate only during particularly demanding tasks (Dembitskaya et al., 2022) - potentially explaining some of the apparently conflicting results.

Increases in MR field strength and improved MRS methodology have enhanced the detection of glutamate and Gamma-Aminobutyric Acid (GABA), the primary excitatory and inhibitory neurotransmitters respectively, presenting compelling targets for fMRS studies. fMRS studies of the primary visual cortex at 7T have shown that prolonged visual stimulation induces increased levels of glutamate and lactate, and reductions in glucose and aspartate (Bednařík et al., 2015; Schaller et al., 2013). Prolonged motor activation has also been studied with fMRS, showing task-induced increases in glutamate and lactate in primary motor cortex (Schaller et al., 2013; Volovyk & Tal, 2020) - findings consistent with prolonged visual stimulation. The MEGA-PRESS MRS sequence (Mescher et al., 1998) is a popular approach for detecting metabolites with weakly J-coupled spins, and may be optimised to measure GABA or lactate by reducing spectral overlap and artefacts from scalp lipids. This approach has been applied to fMRS of the primary motor cortex, detecting an increase in lactate (Koush et al., 2019) and a decrease in GABA (Chen et al., 2017) in response to prolonged activation.

Whilst high field (7 Tesla and above) MRS offers improved SNR and reduced spectral overlap, 3 Tesla MR systems are currently more widely available, and offer a simpler transition from the research setting to clinical use. Despite the popularity of 3 Tesla for neuroimaging applications, vendor-supplied single-voxel MRS implementations are currently based on the PRESS sequence (Bottomley, 1987), which is known to suffer from localisation inaccuracies and signal loss - particularly for lactate (Yablonskiy et al., 1998). The clinical MRS research community has established a consensus on the use of the semi-LASER sequence over PRESS at 3 Tesla to resolve these technical limitations (Deelchand et al., 2018; Oz & Tkáč, 2011; Wilson et al., 2019), however the application of semi-LASER to fMRS remains underexplored, despite its potential advantages.

In this study, we employ short echo-time semi-LASER MRS to measure metabolic changes in the primary motor cortex during a prolonged task at a field strength of 3 Tesla. Compared to conventional PRESS we expect improved estimates of glutamate, due to reduced evolution of strongly J-coupled spins associated with the train of refocusing pulses used in semi-LASER, and improved lactate estimates due to reduced chemical shift displacement. Both these molecules are known to change in response to brain activation, suggesting short echo-time semi-LASER MRS may be a particularly effective acquisition method for fMRS studies at 3 Tesla. In addition to conventional fMRS analysis, whereby dynamic spectra are analysed individually with MRS fitting algorithms (Provencher, 1993; Wilson, 2021a), we introduce a novel alternative based on the application of the General Linear Model (GLM) directly to fMRS spectral time-courses. This “mass-univariate” approach is widely used for fMRI analysis, and is gaining popularity for spectral analysis of electrophysiology data (Quinn et al., 2024).

## 2. Methods

### 2.1. Participants and functional task

A total of 23 (17 female) right-handed participants with a mean age of 21.6 years were recruited for the study. The study was conducted according to the principles expressed in the Declaration of Helsinki and participants gave written informed consent before data collection.

At the start of each session, participants were introduced to the MR environment and instructed on the two tasks to be performed inside the scanner. The primary experiment, to be completed concurrently with fMRS acquisition, was to apply rhythmic clenching to a dynamometer held in the right hand, with the participant’s right arm relaxed and by their side in a supine body position. Participants were instructed to pay attention to a fixation cross for 3 minutes in the first “rest” phase of the experiment. In the following “task” phase, lasting a total of 8 minutes, participants followed visual prompts to apply a rhythmic squeezing force to the dynamometer. Squeezing was instructed at a rate of 1 Hz by repeatedly displaying the words “squeeze” and “relax”, with durations of 0.7 and 0.3 seconds respectively (Chen et al., 2017). Participants were instructed to squeeze at each prompt with approximately half their full strength. Finally, a second “rest” phase, lasting 14 minutes, was completed. The total fMRS experiment lasted 25 minutes.

A secondary experiment was completed concurrently with fMRI acquisition and constituted a 30-second “rest” block, followed by a 30-second “task” block and a final 60-second “rest” block. These data were acquired to ensure the fMRS voxel contained brain tissue hemodynamically responsive to the hand-clenching task. All other aspects of the secondary task were identical to the first.

The MR-compatible dynamometer was combined with a compatible amplifier interface, laptop and associated software (BIOPAC Systems, Inc, CA, USA) for real-time monitoring of the applied force to ensure participants were performing the two experiments correctly. Visual stimuli were presented using Psychopy software (Peirce et al., 2019) version 2023.1 onto a projector screen positioned inside the scanner bore and behind the participant’s head - viewed via a mirror attached to the head-coil. A Propixx system (VPixx Technologies Inc, Canada) was situated outside the scanner room and used to project images onto the screen via a waveguide in line with the scanner bore.

### 2.2. MR acquisition

MR data were acquired with a 3 Tesla Siemens Magnetom Prisma (Siemens Healthcare, Erlangen, Germany) system using a 32-channel receiver head coil array. A T1-weighted MRI scan was acquired sagittally with a 3D-MPRAGE sequence: FOV = 208 × 256 × 256 mm, resolution = 1 × 1 × 1 mm, TE / TR = 2 ms / 2000 ms, inversion time = 880 ms, flip angle = 8°, GRAPPA acceleration factor = 2 (4 min 54 s scan duration).

Single voxel fMRS was performed using the semi-LASER sequence (Deelchand et al., 2018) with a 25 mm-sided cubic excitation region. 750 transients were acquired with a 90 degree flip angle; TE / TR = 28 ms / 2000ms and VAPOR water suppression (25 min scan duration). B_0_ shimming was performed using the “brain” method implemented by the vendor. MRS data was exported and converted to NIfTI MRS format (Clarke et al., 2022) as 2048 complex data points acquired with spectral width of 2000 Hz. The MRS voxel was positioned over the hand motor cortex region in the left hemisphere based on established anatomical landmarks (Caulo et al., 2007) identified from the immediately preceding T1-weighted images (Fig. 1A).

**Fig. 1.**
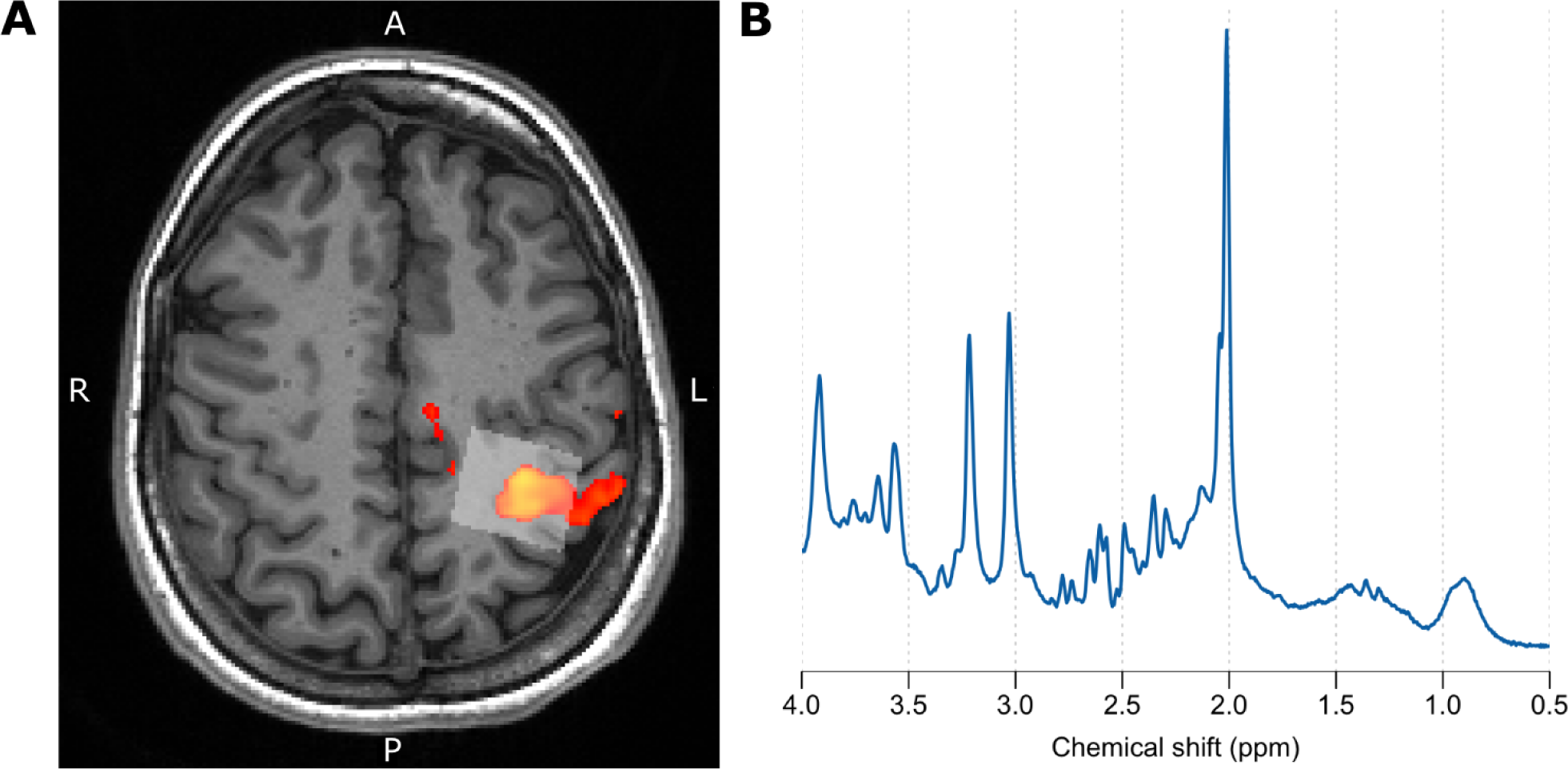
A) An example fMRS voxel (25mm-sided cube in translucent white) placed over hand motor cortex with fMRI BOLD activation overlaid as a colour map. B) mean spectrum averaged over 750 transients following frequency and phase correction.

fMRI data was acquired using a multi-band accelerated GRE-EPI sequence with an isotropic voxel resolution of 2.5 mm over a 210 x 210 x 142.5 mm FOV (AP x LR x FH). 80 fMRI volumes (each containing 57 transverse slices) were acquired with a TR of 1500 ms; TE of 35 ms; 71 degree excitation flip angle; A>>P phase encoding direction and multi-band acceleration factor of 3 (2 min scan duration).

Trigger pulses were generated by the fMRS and fMRI sequences at each TR to ensure temporal synchronisation between the functional tasks and data acquisition. Padding was applied to the inside of the head coil to aid participant comfort and minimise head motion.

### 2.3. Conventional fMRS analysis

Temporal instability in the frequency offset and zero-order phase for each dynamic spectrum was corrected using the RATS method (Wilson, 2019). The 4-1.9 ppm range (containing the strongest metabolite signals) from the mean uncorrected spectrum calculated over the 750 transients for each participant was used as the reference spectrum for correction. The mean corrected spectrum for each participant was then further frequency- and phase-aligned to a simple simulated spectrum composed of three resonances with equal intensity at 2.01, 3.03 and 3.22 ppm with 4 Hz Lorentzian line-broadening applied - using the RATS method. These “participant-level” frequency and phase corrections were then applied back to the individually corrected dynamics to ensure that every single spectrum was consistently phase- and frequency-aligned across all participants. Dynamic spectra from each participant were amplitude normalised to the peak height of the tCr resonance at 3.03 ppm measured from the dynamic mean spectrum on a per-participant basis.

fMRS data quality was assessed by: 1) visual inspection of mean spectra; 2) dynamic spectrograms; 3) spectral SNR based on the peak height of the tNAA resonance and 4) linewidth of the tNAA resonance at 2.01 ppm.

For spectral fitting, corrected spectra were averaged into temporal blocks of 50 spectra to enhance spectral SNR, resulting in 15 dynamic spectra per participant. Conventional spectral fitting was performed for each participant separately, and a further analysis was performed following averaging the 15 dynamic spectra across all participants.

Spectral fitting was performed using a simulated basis containing the following set of standard brain metabolites: alanine (Ala), aspartate (Asp), creatine (Cr), gamma-Aminobutyric acid (GABA), glucose (Glc), glutamine (Gln), glutathione (GSH), glutamine (Gln), glycine (Gly), glycerophosphocholine (GPC), myo-inositol (Ins), lactate (Lac), N-acetylaspartate (NAA), N-acetylaspartylglutamate (NAAG), phosphocholine (PCho), phosphocreatine (PCr), phophosphoethanolamine (PEth), scyllo-inositol (sIns) and taurine (Tau). The commonly used set of simulated broad signals to model macromolecular and lipid signals present in ^1^H MRS data were also included in the basis, see Table 1 of (Wilson et al., 2011) for the full listing.

Spectral fitting was performed initially using the ABfit algorithm (Wilson, 2021a) and repeated using LCModel (Provencher, 1993) to explore consistency of findings across two different methods. Both algorithms were applied with default fitting parameters and an identical basis set as described above. Metabolite levels were divided by the tCr level (sum of PCr and Cr) estimated from the mean spectrum across all time points.

### 2.4. Exploratory fMRS spectro-temporal statistical modelling

The standard approach for fMRS analysis involves temporal averaging of spectra to boost SNR, and therefore the accuracy of metabolite estimates, followed by the use of conventional spectral fitting algorithms to measure dynamic metabolite changes. More recently, we have shown how the application of multiple univariate statistical tests directly to individual MRS data points can support findings from spectral fitting, and potentially reveal novel information on individual differences in neurometabolic profiles (Vella et al., 2023; Wu et al., 2022). A similar approach may be easily adapted to fMRS, where each frequency domain data point can be treated as an independent time course to be fitted with a linear model incorporating the predicted metabolite dynamics. Statistical measures may then be derived from each fit to indicate the likelihood of a given frequency region being associated with a particular functional task. Such an approach is analogous to conventional fMRI analysis - where linear fits incorporating the predicted BOLD response are applied to the time course of each spatially encoded voxel.

Following frequency and phase alignment and normalisation steps described earlier, an additional 2 Hz Gaussian line broadening was applied to enhance spectral SNR. Spectra were cropped to the region between 0 and 4.3 ppm and asymmetric least squares baseline correction was applied (Eilers & Boelens, 2005). Since the true dynamic responses of lactate and glutamate to visual stimuli are yet to be established, a simple boxcar function was assumed, with values of 0 during rest blocks and 1 in the task block. Linear regression of the boxcar function was applied to each spectral time course for participant-averaged data following the preprocessing steps described in section 2.4. Temporal averaging was not performed.

All spectral processing, fitting and associated statistics and plots were performed using the spant MRS analysis package (Wilson, 2021b) implemented in the R statistical programming language (R Core Team, 2021).

### 2.5. fMRI analysis

BOLD activation maps were generated using FEAT, distributed as part of the FSL software package (Woolrich et al., 2009), version 6.0.6.1. A standard analysis pipeline was used, incorporating motion correction and spatial registration to a defaced T1-weighted MPRAGE anatomical scan. 5 mm of spatial smoothing was applied to the EPI data, and a z-threshold of 5.3 was used to generate statistical maps to confirm concordance between MRS voxel placement and BOLD activation.

## 3. Results

### 3.1. fMRS data quality

Manual inspection of spectrograms prior to frequency and phase correction revealed significant movement artefacts for 3 of the 23 participants and these scans were removed from subsequent analysis. An additional scan was removed due to an unusually high degradation in tNAA lineshape FWHM from approximately 0.05 to 0.07 ppm, potentially arising from scanner instability, or gradual participant movement throughout the scan. One participant was unable to complete the fMRI task due to fatigue, however the fMRS data was still included in subsequent analyses since they were able to complete the associated task, as confirmed by real-time observation of the force applied to the dynamometer. The single-shot SNR of each dynamic spectrum was measured and a dynamic median value calculated for each fMRS scan based on the maximum peak height of the tNAA resonance at 2.01 ppm. The mean of these single-shot SNR values was calculated across the 19 good-quality scans as 31 with a standard deviation of 3.4. Similarly, line widths were estimated from the tNAA resonance of the dynamic mean spectrum. The mean linewidth calculated across the 19 good-quality scans was 0.047 ppm with a standard deviation of 0.010 ppm.

### 3.2. Conventional fMRS analysis

Time-courses for participant-averaged spectra are plotted for glutamate and lactate in Fig. 2. Glutamate levels appear consistent throughout the functional task whereas lactate shows a steady increase. A t-test between the metabolite levels during the task vs rest blocks (data points within vs outside the red boundary in Fig. 2) support this observation, with lactate demonstrating a statistically significant change: t(4.7) = 3.98, p = 0.012, in contrast to glutamate: t(11.0) = 0.78, p = 0.45.

**Fig. 2.**
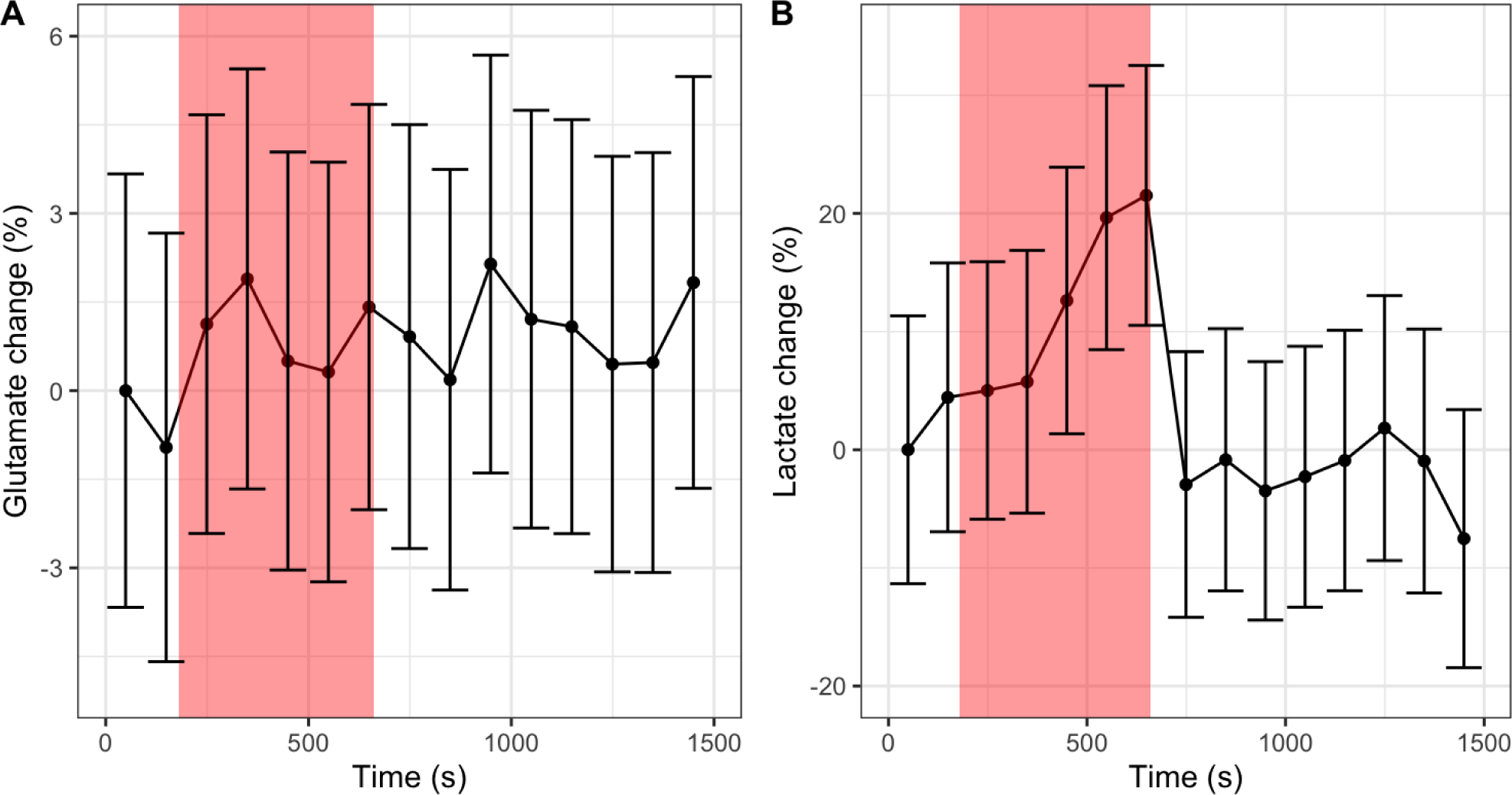
Time-courses for A) glutamate and B) lactate estimated from spectral fitting of participant-averaged spectra using the ABfit method. Error bars represent the standard deviation estimated from Cramer-Rao Lower Bounds. The translucent red region represents the task block.

Fig. 3 shows the mean time-courses for glutamate and lactate calculated over the fitting results derived from each participant separately. Using this alternate analysis approach reveals a change in statistical significance for glutamate (t(7.5) = 3.83, p = 0.0056) and lactate (t(5.7) = 0.62, p = 0.56), between rest and task states, when compared to fitting results from the participant-averaged spectra (Fig. 2).

**Fig. 3.**
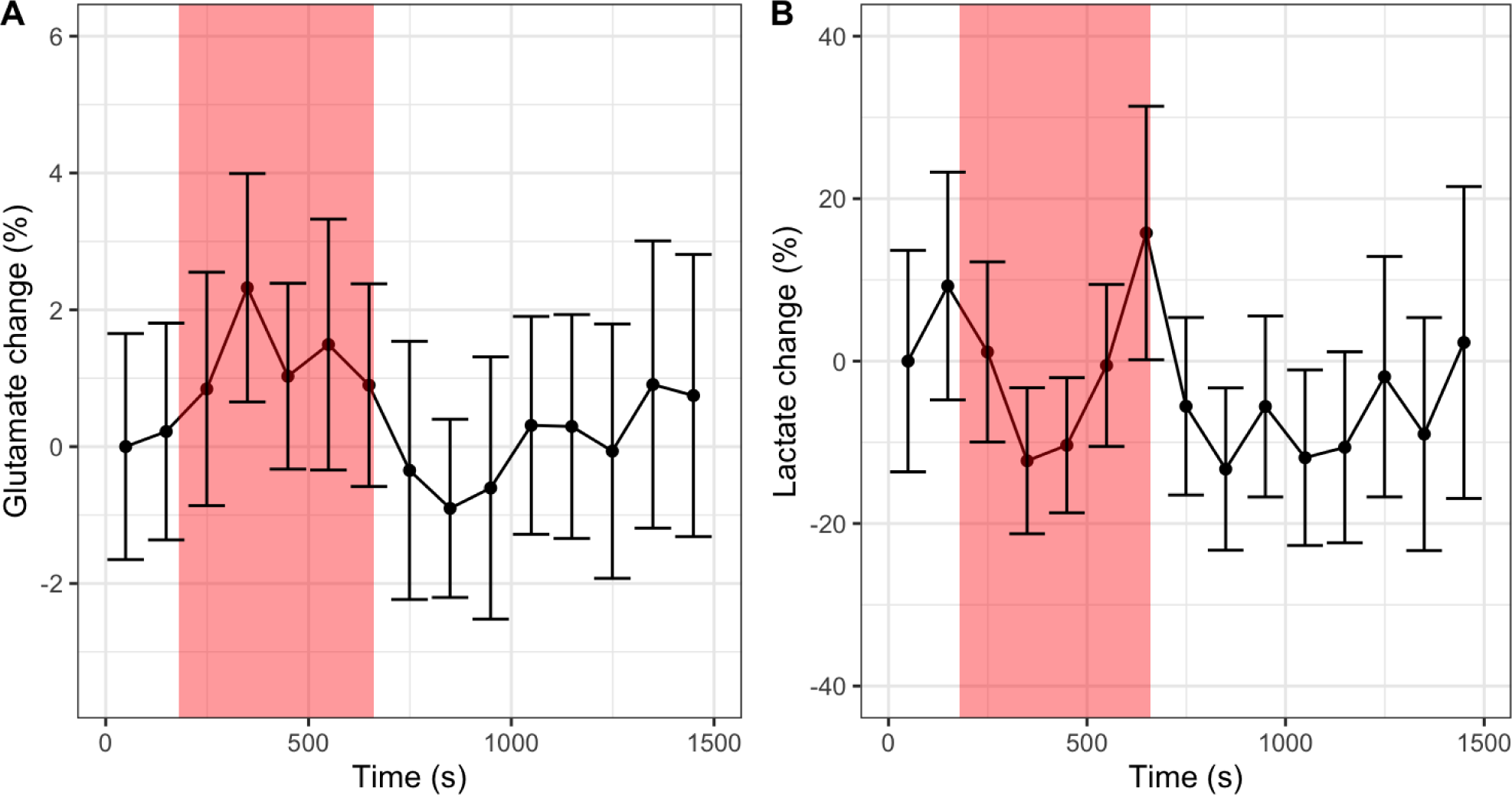
Mean time-courses for A) glutamate and B) lactate estimated from spectral fitting of individual participant spectra using the ABfit method. Error bars represent the standard error across participants. The translucent red region represents the task block.

The discordance between metabolite dynamics estimated from participant mean spectra compared to mean levels from individual participant scans is unexpected, and suggests a level of instability present in the spectral data, the analysis methodology or both. Spectrograms were generated for each individual participant scan and the mean participant scan to explore sources of temporal variance. The dynamic mean spectrum was subtracted from each spectrogram. 4 Hz Gaussian line-broadening and a linear baseline correction was also applied to each spectrum to aid the visualisation of small temporal changes. Fig. 4A shows a typical example of an individual participant spectrogram, where the primary source of temporal variance is evident in the spectral region between 1 and 1.5 ppm. The smooth spectral appearance, combined with the frequency range, strongly suggests this source of variance originates from out-of-volume scalp lipids - a commonly observed artefact and confound for lactate measurement in ^1^H MRS brain data. Fig. 4B shows how this artefact is reduced from approximately ±15% to ±4% the height of the tCr resonance in the participant mean spectrum compared to the individual participant spectrogram in part A. This potentially explains the disagreement between lactate dynamics observed in Fig. 2B compared to Fig. 3B as the effectively random phase of the lipid artefact results in its suppression in the participant-averaged spectra.

**Fig. 4.**
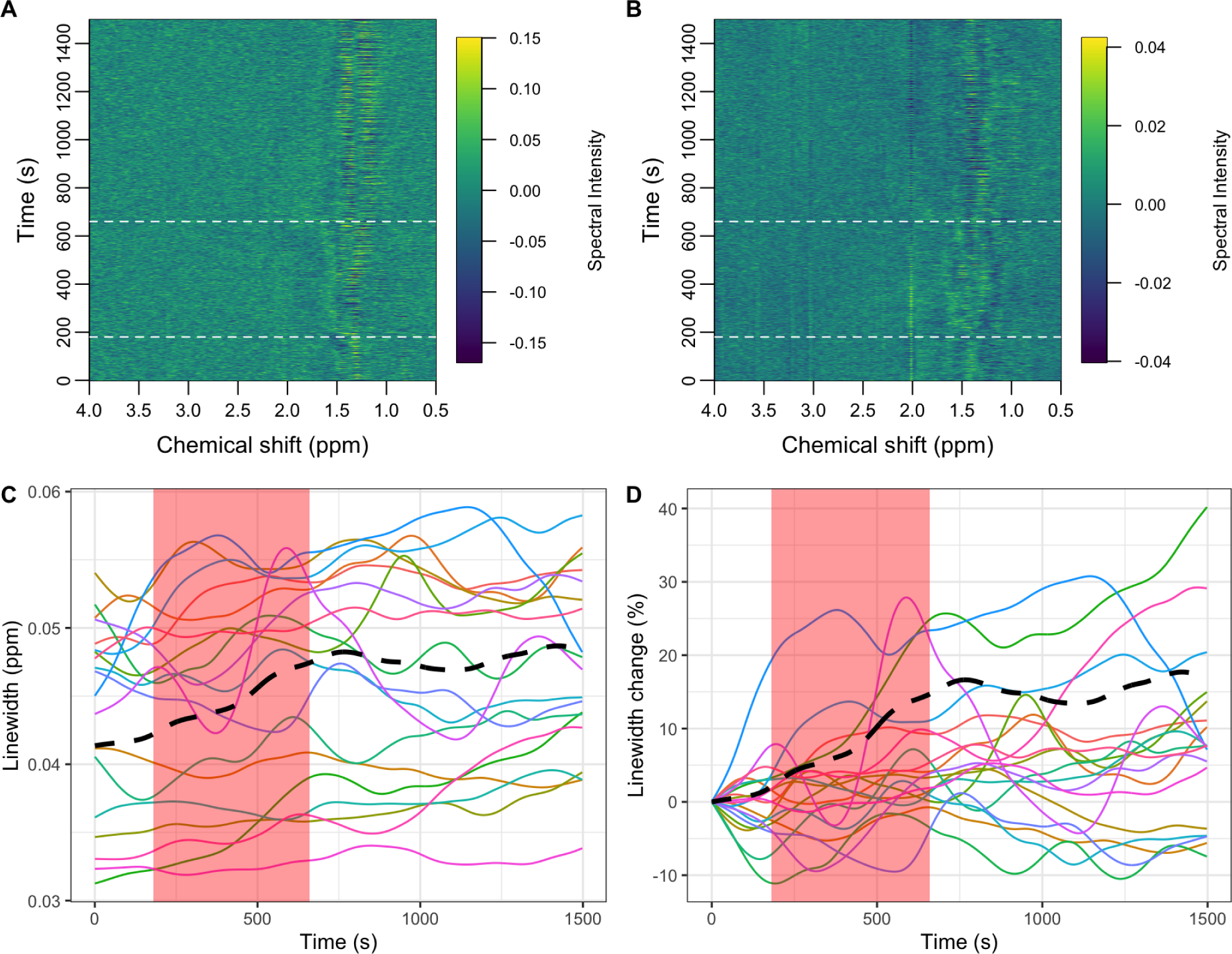
Spectrograms of A) an example participant fMRS scan, B) participant averaged fMRS scan. The dynamic mean spectrum was subtracted from both spectrograms to highlight small temporal variations. Dashed horizontal white lines represent the transitions between rest and task states. C) the smoothed tNAA linewidth for individual participants (coloured lines) and participant averaged scan (black dashed). D) same as C) but measured as a percentage change from the first time-point.

In addition to a reduced lipid artefact, Fig. 4B shows temporal changes in the spectral intensity of the primary singlet resonances of tNAA, tCr and tCho at 2.01, 3.03 and 3.22 ppm respectively. Whilst similar changes in the intensity of these resonances are present in the individual participant data, the improved SNR of the average participant spectra aids visualisation of this type of variability. These changes are attributed to dynamic linewidth variability, potentially resulting from a slow degradation in magnetic field homogeneity, with Fig. 4 part D showing gradual lineshape increases of up to 40% in the worst case. The participant average data (dashed black line in parts C and D) however shows a milder increase of up to 20% (from 0.041 to 0.049 ppm). Spectral fitting approaches have been shown previously to be sensitive to changes in linewidth (Mangia et al., 2006; Near et al., 2013), therefore we propose that differences in SNR and linewidth variability between the individual subject analysis (Fig. 3A) and subject averaged analysis (Fig. 2A) may explain the differences in estimated glutamate dynamics.

Exploratory analysis was also performed with the LCModel fitting algorithm (Provencher, 1993), since this fitting method is more commonly used for fMRS analysis. Time-courses for glutamate and lactate, measured from the participant mean spectra, are shown in Fig. S1. Applying a t-test between the metabolite levels during the task vs rest blocks gave statistically significant differences for both glutamate (t(10.4) = 2.65, p = 0.023) and lactate (t(4.6) = 3.79, p = 0.015). Consistent results were also found for the analysis of mean time courses from individual participant scans, with statistically significant increases in glutamate (t(10.1) = 2.90, p = 0.016) and lactate (t(5.1) = 3.41, p = 0.018) during the task block (Fig. S2).

### 3.3. Exploratory fMRS spectro-temporal statistical modelling

Further exploratory analysis of the participant mean spectral data was performed using a novel processing approach performed directly on the spectral data points, obviating the need for a simulated basis set or complex fitting algorithm. Fig. 5A highlights the spectral regions most different between the task and rest states by fitting each spectral time-course with a simple boxcar function corresponding to the task state. In this preliminary analysis, the spectral regions most strongly associated with the task are the three primary singlet resonances of tNAA, tCr and tCho and the strong myo-inositol multiplet at 3.55 ppm. These differences are due to the drift in linewidth (see Fig. 4) rather than genuine metabolic changes, therefore we added a nuisance regressor to the boxcar function which was calculated from the temporally smoothed time-course of the integrated spectral region between 1.97 to 2.04 ppm. Fig. 5B shows how the incorporation of the nuisance regressor significantly reduces the intensity of the spectral regions most associated with lineshape drift. The primary resonances of lactate at 1.28 and 1.35 ppm are present in both the basic analysis (Fig. 5A) and the analysis including the nuisance component (Fig. 5B) suggesting the lactate changes are not strongly influenced by lineshape changes. Conversely, the primary glutamate resonance at 2.35 ppm shows only a weak statistical association to the task compared to lactate in both analyses.

**Fig. 5.**
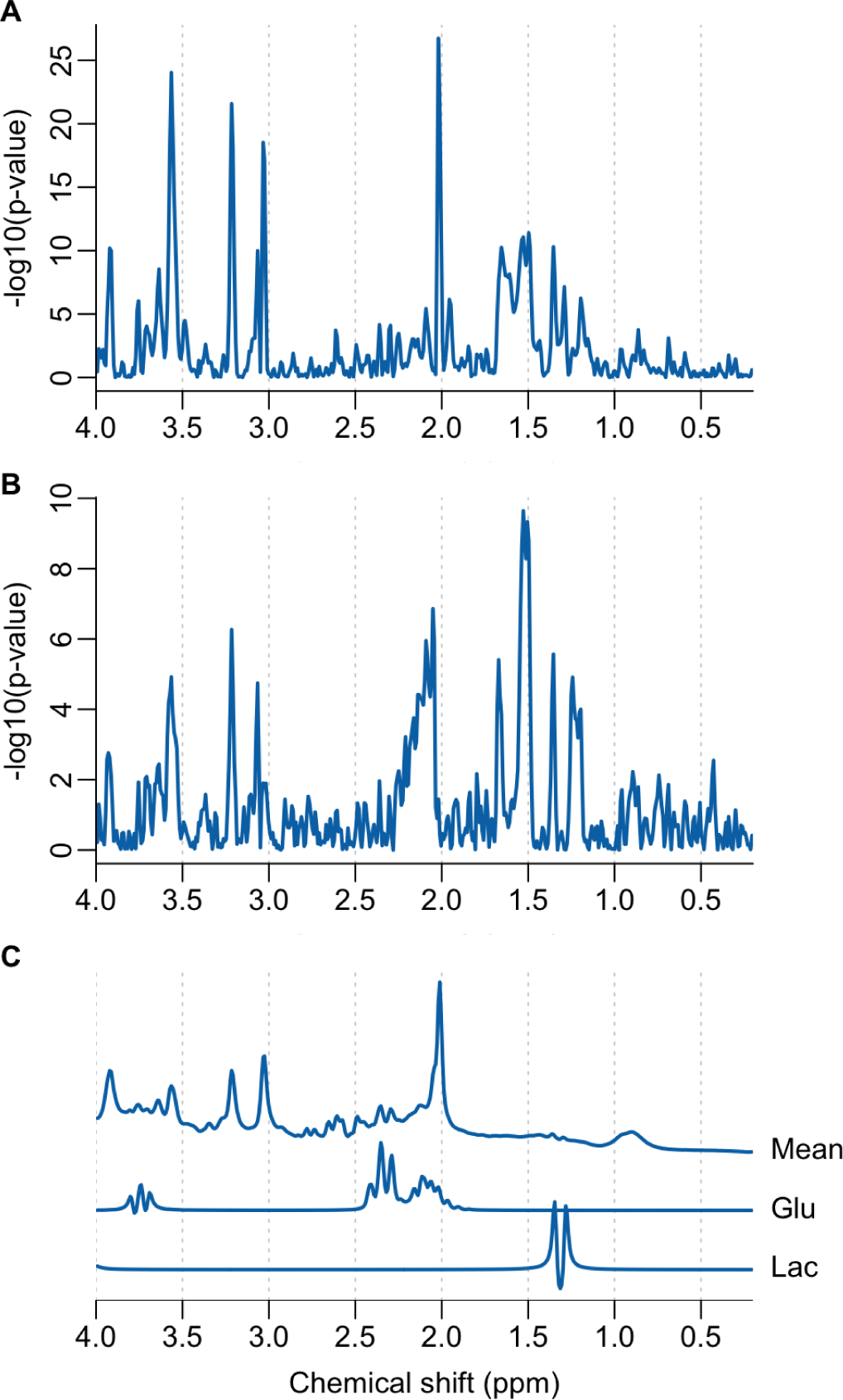
A) A Manhattan plot illustrating spectral regions temporally associated with the functional task, assuming a simple boxcar function. B) Same as A) but with an added nuisance regressor to suppress the influence of lineshape variability. C) mean fMRS spectrum with simulated glutamate (Glu) and lactate (Lac) signals. Simulated signals are scaled to have similar maximum intensities to aid the assignment of parts A) and B).

Metabolite dynamics in longer block designs have been shown to have a lagged response to tasks (Bednařík et al., 2015), with some metabolites taking up to 2 minutes to plateau followed by a slow decline once the task has stopped. Fig. 6 shows how delaying the boxcar function by 2 minutes further suppresses the primary singlet spectral regions most sensitive to lineshape changes, whilst maintaining a strong association with lactate levels. Very little association with the primary resonance of glutamate is apparent with the lagged response function, however we do note a weak association with potential resonances at 2.74 and 2.78 ppm which we tentatively assign to aspartate. A -5.4% decrease in aspartate in response to visual stimulation has been reported previously (Bednařík et al., 2015), therefore we chose to examine these levels estimated from conventional spectral fitting. Fig. S3 shows the aspartate levels estimated from the participant-averaged spectra. An approximately 4% decrease is seen at the 5th time point in agreement with (Bednařík et al., 2015). Statistical testing between the rest and task time points for aspartate did not reach statistical significance (t(6.9) = -1.78, p = 0.11), however delaying the task window by one time point (100 seconds), similar to Fig. 6, did reveal a statistically significant difference (t(12.1) = -4.33, p = 0.00097). The delayed boxcar regressor weighings (beta weights) are shown in Fig. S4A and confirm a task-related increase in lactate (positive beta weightings at 1.28 and 1.35 ppm) and decrease in aspartate (negative beta weightings at 2.74 and 2.78 ppm).

**Fig. 6.**
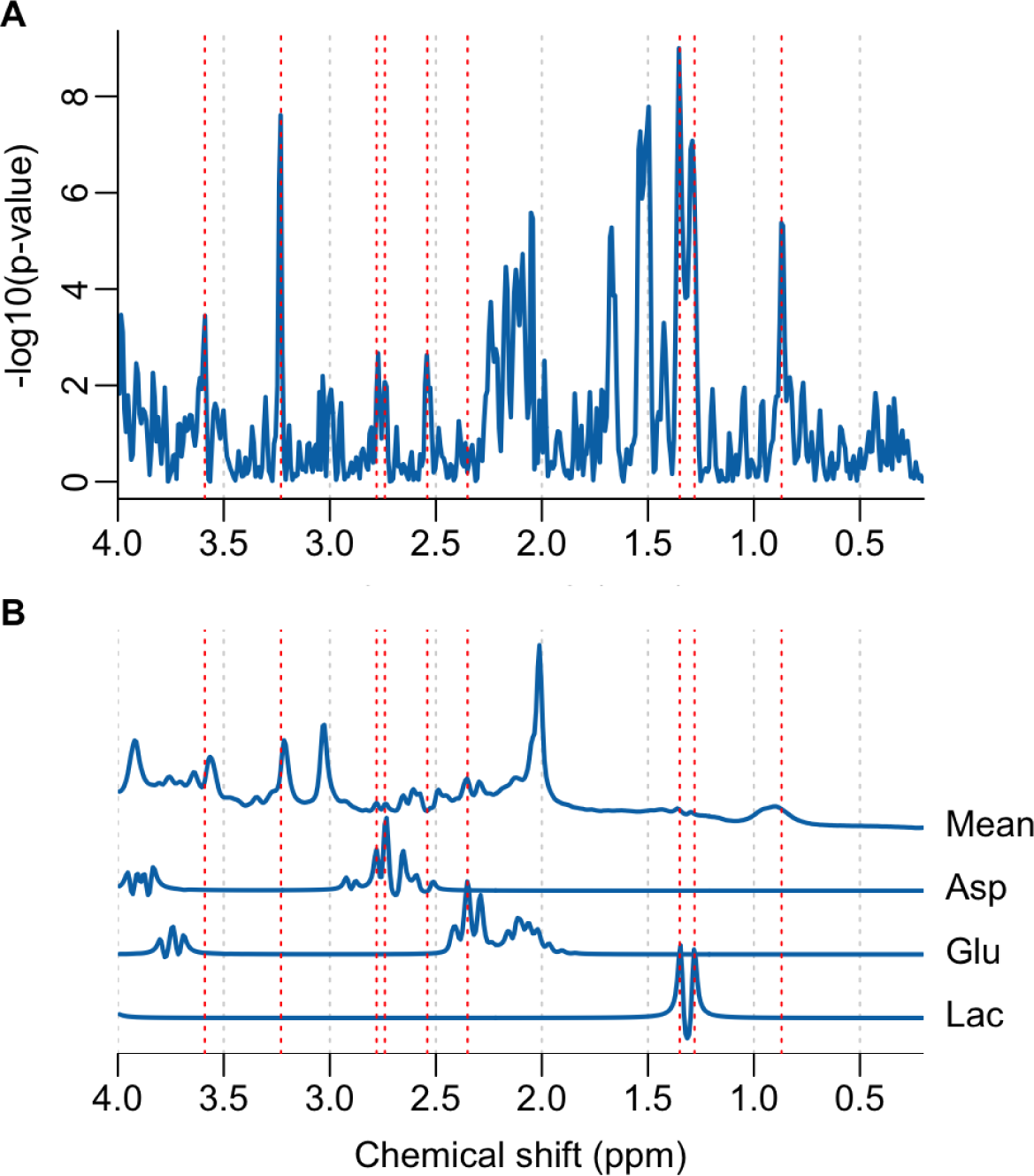
A) A Manhattan plot illustrating spectral regions temporally associated with the functional task, assuming a simple boxcar function delayed by 120 seconds. B) mean fMRS spectrum with simulated glutamate (Glu), lactate (Lac) and aspartate (Asp) signals. Simulated signals are scaled to have similar maximum intensities to aid the assignment of part A). Dashed red lines highlight spectral features at 0.87, 1.28, 1.35, 2.35, 2.54, 2.74, 2.78, 3.23 and 3.59 ppm.

## 4. Discussion

Previous studies of prolonged motor activation have reported increases in glutamate and lactate in the motor cortex and our findings largely support these observations (Chen et al., 2017; Koush et al., 2019; Schaller et al., 2013; Volovyk & Tal, 2020). However, we also observed how measures of statistical significance were sensitive to the analysis approach, showing differences depending on how participant-averaged results were compiled and the choice of spectral fitting algorithm. Inspection of spectral variability (Fig. 4) demonstrated that dynamic changes in metabolite linewidth and lipid artefacts were the primary sources of variance and that averaging across participant spectra may reduce these artefacts, potentially explaining some of the inconsistencies between analysis approaches (Fig. 2 vs 3). Differing results between spectral fitting strategies have been reported previously (Marjańska et al., 2022), and are indicative of the inherent instabilities of non-linear analyses in the presence of noise. Whilst fitting stability may be imposed through various forms of regularisation (e.g. Bayesian priors), this ultimately results in some degree of bias due to the “bias-variance tradeoff”. Even within the same fitting approach, variation in results may arise from minor changes in basis set constituents (Demler et al., 2024), further emphasising the inherent instability of spectral fitting. For large metabolite changes, typically seen with clinical MRS, these instabilities are less problematic; however changes are typically on the order of less than 10 percent for ^1^H fMRS and therefore more stable analysis approaches are required.

In this study, we present an alternate analysis approach, based on the GLM, to improve the stability of fMRS findings. In contrast to conventional spectral fitting, no assumptions about the basis-set are made, avoiding one potential source of variability between analyses (Demler et al., 2024). Simple linear modelling is performed directly on spectral data points using the “mass univariate” approach, which is well-established for fMRI, and is becoming a more popular tool for electrophysiology (Quinn et al., 2024) and conventional MRS (Vella et al., 2023; Wu et al., 2022). While further validation of the approach is required, we show here how it may be used to support or question findings from conventional analyses. The corroboration between conventional fitting, which shows an approximately 20% increase in lactate from baseline levels, and the clear doublet appearance at 1.28 and 1.35 ppm in the Manhattan plot (Fig. 6A) adds confidence to these results. A similar increase in lactate in response to a prolonged motor task (17%) was reported at a field strength of 7 Tesla (Schaller et al., 2014) further supports our findings. This is the first study to show lactate dynamics of prolonged motor activation may be observed at 3 Tesla. Another similar study at 3 Tesla was unable to measure these changes due to unreliable quantitation (Volovyk & Tal, 2020). Possible explanations for this difference in findings may be due to alternate acquisition methodologies (semi-LASER vs PR-STRESS) and differences in voxel dimensions (15.6 ml vs 6 ml). Whilst larger voxel dimensions are expected to be more susceptible to scalp lipid contamination, this may be offset by improved SNR. Our results may also suggest that the spatial extent of the lactate response extends beyond a 6 ml volume, since unaffected tissue would effectively “dilute” the measured percentage change, yet we still observe a relatively large change of 20% - consistent with 7 Tesla studies.

Glutamate changes of approximately 2%, as measured from spectral fitting (Fig. 3A), were comparatively smaller than the values of around 4% reported elsewhere at 3 Tesla (Volovyk & Tal, 2020). We also found inconsistency between the analysis of participant average spectra (Fig. 2A), the participant-averaged metabolite levels (Fig. 3B) and the GLM spectral modelling (Fig. 6A). Whilst multiple resonances around 2.2 ppm in Fig. 6A are consistent with glutamate (Fig. 6B), the strongest resonances expected around 2.35 ppm are absent. This may imply that fitting differences are driven by changes in the unassigned resonances at 2.2 ppm, rather than glutamate, or that lineshape variance is correlated with glutamate changes, obscuring detection accuracy. We also note that our voxel volume of 15.6 ml was larger than in similar studies (∼10 ml), which may suggest that glutamate changes have a smaller spatial extent than lactate.

The exploratory spectral GLM analysis (Fig. 6A) highlighted a potential metabolic response in resonances at 2.74 and 2.78 ppm. These frequencies are in good agreement with those expected from aspartate, prompting further exploratory analysis of the conventionally fitted data. An approximately 4% decrease in aspartate is shown in Fig. S3, consistent in magnitude and direction to changes observed during extended visual stimulation at 7 Tesla (Bednařík et al., 2015). Whilst exploratory in nature, the agreement between spectral GLM, conventional analysis and high field studies suggest the change is likely genuine and demonstrates how novel analysis approaches have the potential to improve the sensitivity and reliability of fMRS.

In addition to those already mentioned, Fig. 6A shows a number of other spectral regions with a statistical dependence on the functional task. The peak at 3.23 ppm is assigned to resonances from choline, phosphocholine and glycerophosphocholine and previous work has shown alterations in choline levels during reversal learning (Bell et al., 2018). Whilst this resonance is also known to be sensitive to lineshape alterations (Fig. 5A), the absence of a similar artefact in the creatine region suggests a change in concentration or frequency shift of the choline resonances may be genuine, and warrants further study. Further resonances at 0.87, 1.50, 1.68, 2.54 and 3.59 ppm cannot be assigned with confidence, however we note some similarity to resonances expected from branched-chain amino acids.

Alternative MRS analysis approaches involve 2D fitting, whereby multiple spectra are fit to a metabolite basis simultaneously - reducing the number of parameters requiring optimisation resulting in more accurate estimates. Whilst the advantages of 2D fitting in MRS have been long established in various contexts (Chong et al., 2011; Schulte & Boesiger, 2006; Van Ormondt et al., 1990) there has been renewed interest in the application of 2D fitting to functional and diffusion MRS (Clarke et al., 2024; Tal, 2023). Other approaches involve combining conventional 1D fitting with a GLM applied to the metabolite time-courses (Ligneul et al., 2021). How these methods compare to the direct linear modelling of spectral time-courses introduced here, particularly in the presence of confounding temporal instabilities, presents a key area for the future development of fMRS analysis methodology.

## Conclusions

We have shown that lactate dynamics in response to a prolonged motor task are observed using short-echo time semi-LASER at 3 Tesla. A novel approach for fMRS data analysis is introduced, based on direct linear modelling of spectral time-courses, demonstrating potential for measuring small changes associated with aspartate and other unassigned molecules. Future work includes reducing the effect of spectral confounds, such as scalp lipid contamination and temporal lineshape variability, and further validation of our new analysis approach using experimentally acquired and simulated datasets.

## Data and Code Availability Statement

All code to generate the results presented in this paper is available from: https://github.com/martin3141/fmrs_motor_paper

All data to generate the results presented in this paper are available from: https://doi.org/10.5281/zenodo.11190359

## Author Contributions

Conceptualization (M.M., A.J.Q., P.G.M., M.A.J.A., M.W.); Methodology (M.M., A.J.Q, P.G.M., M.A.J.A., M.W.); Software (M.W., A.J.Q); Formal analysis (M.M., M.W.); Investigation (M.M., M.W.); Resources (D.K.D., M.A.J.A.); Writing - Original Draft (M.M., M.W.); Writing - Review & Editing (M.M., K.D., D.K.D., A.J.Q., P.G.M., M.A.J.A., M.W.); Supervision (M.A.J.A., M.W.).

## Declaration of Competing Interest

None.

## Acknowledgments

DKD acknowledges funding support from NIH grants NIBIB P41 EB027061 and P30 NS076408. M.A.J.A. was funded by a Biosciences and Biotechnology Research Council (BBSRC) Future Leader Fellowship (BB/M013596/1) and a BBSRC David Phillips Fellowship (BB/R010668/1).

## Notes

### Competing Interest Statement

The authors have declared no competing interest.

## References

Bednařík, P., Tkáč, I., Giove, F., DiNuzzo, M., Deelchand, D. K., Emir, U. E., Eberly, L. E., & Mangia, S. (2015). Neurochemical and BOLD responses during neuronal activation measured in the human visual cortex at 7 Tesla. Journal of Cerebral Blood Flow and Metabolism: Official Journal of the International Society of Cerebral Blood Flow and Metabolism, 35(4), 601–610. 10.1038/jcbfm.2014.233

Bell, T., Lindner, M., Mullins, P. G., & Christakou, A. (2018). Functional neurochemical imaging of the human striatal cholinergic system during reversal learning. The European Journal of Neuroscience, 47(10), 1184–1193. 10.1111/ejn.13803

Bottomley, P. A. (1987). Spatial localization in NMR spectroscopy in vivo. Annals of the New York Academy of Sciences, 508, 333–348. 10.1111/j.1749-6632.1987.tb32915.x

Caulo, M., Briganti, C., Mattei, P. A., Perfetti, B., Ferretti, A., Romani, G. L., Tartaro, A., & Colosimo, C. (2007). New Morphologic Variants of the Hand Motor Cortex as Seen with MR Imaging in a Large Study Population. American Journal of Neuroradiology, 28(8), 1480–1485. 10.3174/ajnr.A0597

Chen, C., Sigurdsson, H. P., Auer, D. P., Morris, P. G., Morgan, P. S., Gowland, P. A., & Jackson, S. R. (2017). Activation induced changes in GABA: Functional MRS at 7T with MEGA-sLASER. NeuroImage, 156, 207–213. 10.1016/j.neuroimage.2017.05.044

Chong, D. G. Q., Kreis, R., Bolliger, C. S., Boesch, C., & Slotboom, J. (2011). Two-dimensional linear-combination model fitting of magnetic resonance spectra to define the macromolecule baseline using FiTAID, a Fitting Tool for Arrays of Interrelated Datasets. Magnetic Resonance Materials in Physics, Biology and Medicine, 24(3), 147–164. 10.1007/s10334-011-0246-y

Clarke, W. T., Bell, T. K., Emir, U. E., Mikkelsen, M., Oeltzschner, G., Shamaei, A., Soher, B. J., & Wilson, M. (2022). NIfTI-MRS: A standard data format for magnetic resonance spectroscopy. Magnetic Resonance in Medicine. 10.1002/mrm.29418

Clarke, W. T., Ligneul, C., Cottaar, M., Ip, I. B., & Jbabdi, S. (2024). Universal dynamic fitting of magnetic resonance spectroscopy. Magnetic Resonance in Medicine, 91(6), 2229–2246. 10.1002/mrm.30001

Deelchand, D. K., Kantarci, K., & Öz, G. (2018). Improved localization, spectral quality, and repeatability with advanced MRS methodology in the clinical setting. Magnetic Resonance in Medicine, 79(3), 1241–1250. 10.1002/mrm.26788

Dembitskaya, Y., Piette, C., Perez, S., Berry, H., Magistretti, P. J., & Venance, L. (2022). Lactate supply overtakes glucose when neural computational and cognitive loads scale up. Proceedings of the National Academy of Sciences, 119(47), e2212004119. 10.1073/pnas.2212004119

Demler, V. F., Sterner, E. F., Wilson, M., Zimmer, C., & Knolle, F. (2024). The impact of spectral basis set composition on estimated levels of cingulate glutamate and its associations with different personality traits. BMC Psychiatry, 24(1), 320. 10.1186/s12888-024-05646-x

Díaz-García, C. M., Mongeon, R., Lahmann, C., Koveal, D., Zucker, H., & Yellen, G. (2017). Neuronal stimulation triggers neuronal glycolysis and not lactate uptake. Cell Metabolism, 26(2), 361–374.e4. 10.1016/j.cmet.2017.06.021

Eilers, P. H., & Boelens, H. F. (2005). Baseline correction with asymmetric least squares smoothing. Leiden University Medical Centre Report, 1(1), 5.

Koush, Y., De Graaf, R. A., Jiang, L., Rothman, D. L., & Hyder, F. (2019). Functional MRS with J-edited lactate in human motor cortex at 4 T. NeuroImage, 184, 101–108. 10.1016/j.neuroimage.2018.09.008

Ligneul, C., Fernandes, F. F., & Shemesh, N. (2021). High temporal resolution functional magnetic resonance spectroscopy in the mouse upon visual stimulation. NeuroImage, 234, 117973. 10.1016/j.neuroimage.2021.117973

Maddock, R. J., Buonocore, M. H., Lavoie, S. P., Copeland, L. E., Kile, S. J., Richards, A. L., & Ryan, J. M. (2006). Brain lactate responses during visual stimulation in fasting and hyperglycemic subjects: A proton magnetic resonance spectroscopy study at 1.5 Tesla. Psychiatry Research, 148(1), 47–54. 10.1016/j.pscychresns.2006.02.004

Magistretti, P. J., & Allaman, I. (2018). Lactate in the brain: From metabolic end-product to signalling molecule. Nature Reviews Neuroscience, 19(4), Article 4. 10.1038/nrn.2018.19

Mangia, S., Tkáč, I., Gruetter, R., Van De Moortele, P.-F., Giove, F., Maraviglia, B., & Uğurbil, K. (2006). Sensitivity of single-voxel 1H-MRS in investigating the metabolism of the activated human visual cortex at 7 T. Magnetic Resonance Imaging, 24(4), 343–348. 10.1016/j.mri.2005.12.023

Mangia, S., Tkác, I., Gruetter, R., Van de Moortele, P.-F., Maraviglia, B., & Uğurbil, K. (2007). Sustained neuronal activation raises oxidative metabolism to a new steady-state level: Evidence from 1H NMR spectroscopy in the human visual cortex. Journal of Cerebral Blood Flow and Metabolism: Official Journal of the International Society of Cerebral Blood Flow and Metabolism, 27(5), 1055–1063. 10.1038/sj.jcbfm.9600401

Mangia, S., Tkác, I., Logothetis, N. K., Gruetter, R., Van de Moortele, P.-F., & Uğurbil, K. (2007). Dynamics of lactate concentration and blood oxygen level-dependent effect in the human visual cortex during repeated identical stimuli. Journal of Neuroscience Research, 85(15), 3340–3346. 10.1002/jnr.21371

Marjańska, M., Deelchand, D. K., Kreis, R., & 2016 ISMRM MRS Study Group Fitting Challenge Team. (2022). Results and interpretation of a fitting challenge for MR spectroscopy set up by the MRS study group of ISMRM. Magnetic Resonance in Medicine, 87(1), 11–32. 10.1002/mrm.28942

Mescher, M., Merkle, H., Kirsch, J., Garwood, M., & Gruetter, R. (1998). Simultaneous in vivo spectral editing and water suppression. NMR in Biomedicine, 11(6), 266–272. 10.1002/(sici)1099-1492(199810)11:6<266::aid-nbm530>3.0.co;2-j

Mullins, P. G. (2018). Towards a theory of functional magnetic resonance spectroscopy (fMRS): A meta-analysis and discussion of using MRS to measure changes in neurotransmitters in real time. Scandinavian Journal of Psychology, 59(1), 91–103. 10.1111/sjop.12411

Near, J., Andersson, J., Maron, E., Mekle, R., Gruetter, R., Cowen, P., & Jezzard, P. (2013). Unedited in vivo detection and quantification of γ-aminobutyric acid in the occipital cortex using short-TE MRS at 3 T. NMR in Biomedicine, 26(11), 1353–1362. 10.1002/nbm.2960

Oz, G., & Tkáč, I. (2011). Short-echo, single-shot, full-intensity proton magnetic resonance spectroscopy for neurochemical profiling at 4 T: Validation in the cerebellum and brainstem. Magnetic Resonance in Medicine, 65(4), 901–910. 10.1002/mrm.22708

Prichard, J., Rothman, D., Novotny, E., Petroff, O., Kuwabara, T., Avison, M., Howseman, A., Hanstock, C., & Shulman, R. (1991). Lactate rise detected by 1H NMR in human visual cortex during physiologic stimulation. Proceedings of the National Academy of Sciences, 88(13), 5829–5831. 10.1073/pnas.88.13.5829

Provencher, S. W. (1993). Estimation of metabolite concentrations from localized in vivo proton NMR spectra. Magnetic Resonance in Medicine, 30(6), 672–679. 10.1002/mrm.1910300604

Quinn, A. J., Atkinson, L. Z., Gohil, C., Kohl, O., Pitt, J., Zich, C., Nobre, A. C., & Woolrich, M. W. (2024). The GLM-spectrum: A multilevel framework for spectrum analysis with covariate and confound modelling. Imaging Neuroscience, 2, 1–26. 10.1162/imag_a_00082

R Core Team. (2021). R: A Language and Environment for Statistical Computing. https://www.R-project.org/

Sappey-Marinier, D., Calabrese, G., Fein, G., Hugg, J. W., Biggins, C., & Weiner, M. W. (1992). Effect of Photic Stimulation on Human Visual Cortex Lactate and Phosphates Using 1H and 31P Magnetic Resonance Spectroscopy. Journal of Cerebral Blood Flow & Metabolism, 12(4), 584–592. 10.1038/jcbfm.1992.82

Schaller, B., Mekle, R., Xin, L., Kunz, N., & Gruetter, R. (2013). Net increase of lactate and glutamate concentration in activated human visual cortex detected with magnetic resonance spectroscopy at 7 tesla. Journal of Neuroscience Research, 91(8), 1076–1083. 10.1002/jnr.23194

Schaller, B., Xin, L., O’Brien, K., Magill, A. W., & Gruetter, R. (2014). Are glutamate and lactate increases ubiquitous to physiological activation? A 1H functional MR spectroscopy study during motor activation in human brain at 7Tesla. NeuroImage, 93, 138–145. 10.1016/j.neuroimage.2014.02.016

Schulte, R. F., & Boesiger, P. (2006). ProFit: Two-dimensional prior-knowledge fitting of J-resolved spectra. NMR in Biomedicine, 19(2), 255–263. 10.1002/nbm.1026

Stanley, J. A., & Raz, N. (2018). Functional Magnetic Resonance Spectroscopy: The “New” MRS for Cognitive Neuroscience and Psychiatry Research. Frontiers in Psychiatry, 9. https://www.frontiersin.org/journals/psychiatry/articles/10.3389/fpsyt.2018.00076

Tal, A. (2023). The future is 2D: Spectral-temporal fitting of dynamic MRS data provides exponential gains in precision over conventional approaches. Magnetic Resonance in Medicine, 89(2), 499–507. 10.1002/mrm.29456

Van Ormondt, D., De Beer, R., Mariën, A. J. H., Den Hollander, J. A., Luyten, P. R., & Vermeulen, J. W. A. H. (1990). 2D approach to quantitation of inversion-recovery data. Journal of Magnetic Resonance (1969), 88(3), 652–659. 10.1016/0022-2364(90)90298-N

Vella, O., Bagshaw, A. P., & Wilson, M. (2023). SLIPMAT: A pipeline for extracting tissue-specific spectral profiles from 1H MR spectroscopic imaging data. NeuroImage, 277, 120235. 10.1016/j.neuroimage.2023.120235

Volovyk, O., & Tal, A. (2020). Increased Glutamate concentrations during prolonged motor activation as measured using functional Magnetic Resonance Spectroscopy at 3T. NeuroImage, 223, 117338. 10.1016/j.neuroimage.2020.117338

Wilson, M. (2019). Robust retrospective frequency and phase correction for single-voxel MR spectroscopy. Magnetic Resonance in Medicine, 81(5), 2878–2886. 10.1002/mrm.27605

Wilson, M. (2021a). Adaptive baseline fitting for 1H MR spectroscopy analysis. Magnetic Resonance in Medicine, 85(1), 13–29. 10.1002/mrm.28385

Wilson, M. (2021b). spant: An R package for magnetic resonance spectroscopy analysis. Journal of Open Source Software, 6(67), 3646. 10.21105/joss.03646

Wilson, M., Andronesi, O., Barker, P. B., Bartha, R., Bizzi, A., Bolan, P. J., Brindle, K. M., Choi, I.-Y., Cudalbu, C., Dydak, U., Emir, U. E., Gonzalez, R. G., Gruber, S., Gruetter, R., Gupta, R. K., Heerschap, A., Henning, A., Hetherington, H. P., Huppi, P. S., … Howe, F. A. (2019). Methodological consensus on clinical proton MRS of the brain: Review and recommendations. Magnetic Resonance in Medicine, 82(2), 527–550. 10.1002/mrm.27742

Wilson, M., Reynolds, G., Kauppinen, R. A., Arvanitis, T. N., & Peet, A. C. (2011). A constrained least-squares approach to the automated quantitation of in vivo ^1^H magnetic resonance spectroscopy data. Magnetic Resonance in Medicine, 65(1), 1–12. 10.1002/mrm.22579

Woolrich, M. W., Jbabdi, S., Patenaude, B., Chappell, M., Makni, S., Behrens, T., Beckmann, C., Jenkinson, M., & Smith, S. M. (2009). Bayesian analysis of neuroimaging data in FSL. NeuroImage, 45(1 Suppl), S173-186. 10.1016/j.neuroimage.2008.10.055

Wu, B., Bagshaw, A. P., Hickey, C., Kühn, S., & Wilson, M. (2022). Evidence for distinct neuro-metabolic phenotypes in humans. NeuroImage, 249, 118902. 10.1016/j.neuroimage.2022.118902

Yablonskiy, D. A., Neil, J. J., Raichle, M. E., & Ackerman, J. J. (1998). Homonuclear J coupling effects in volume localized NMR spectroscopy: Pitfalls and solutions. Magnetic Resonance in Medicine, 39(2), 169–178. 10.1002/mrm.1910390202

