## Supplementary material for "Functional Magnetic Resonance Spectroscopy of Prolonged Motor Activation using Conventional and Spectral GLM Analyses": Fig. S

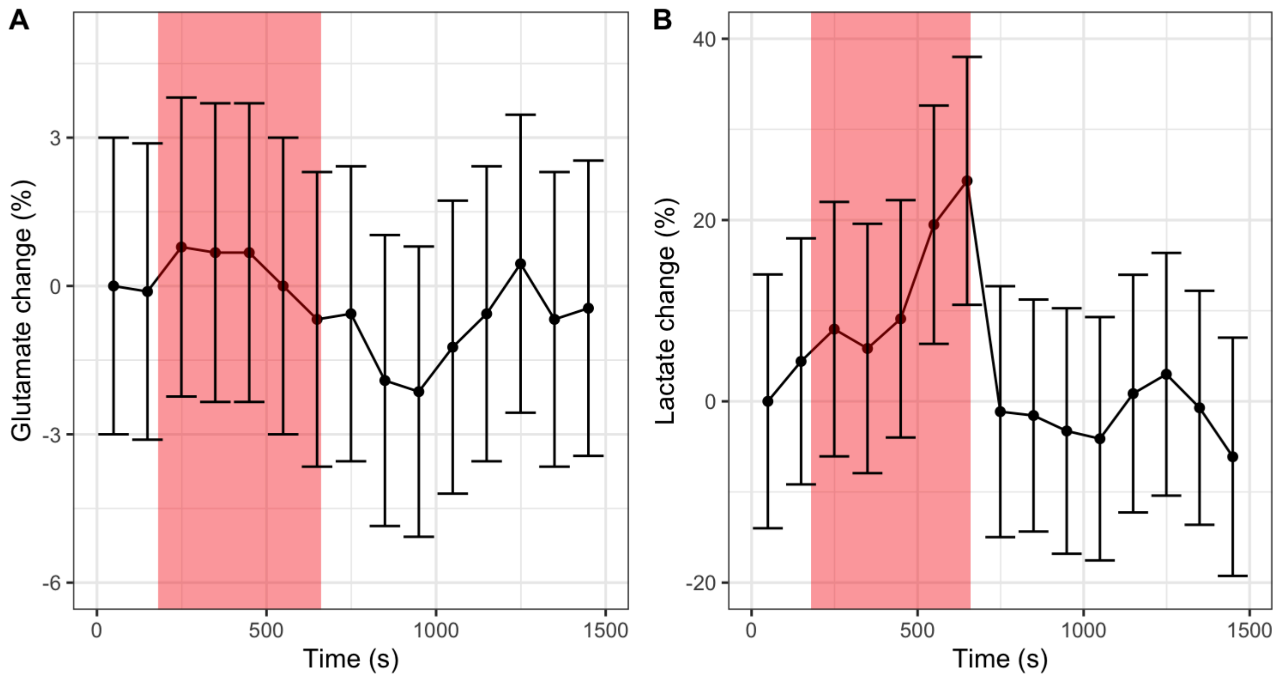


*Fig. S1 Time-courses for A) glutamate and B) lactate estimated from spectral fitting of participant averaged spectra using the LCModel method. Error bars represent the standard deviation estimated from Cramer-Rao Lower Bounds. The translucent red region represents the task block.*

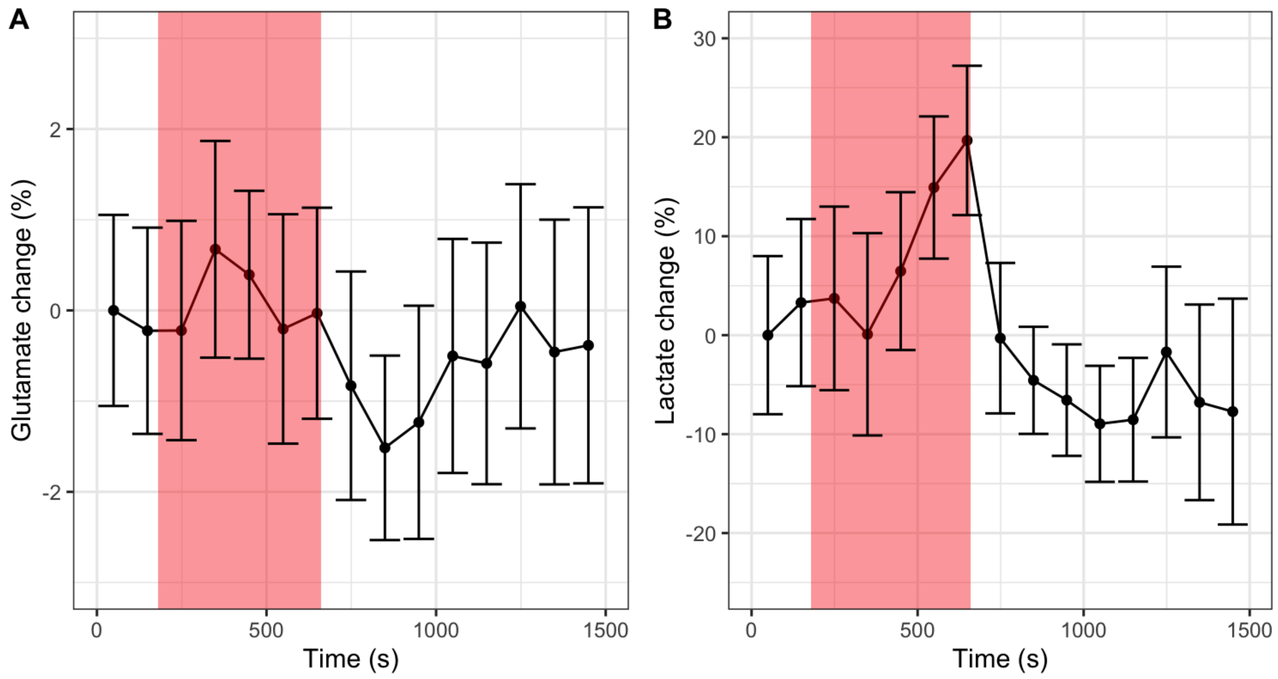


*Fig. S2 Mean time-courses for A) glutamate and B) lactate estimated from spectral fitting of individual participant spectra using the LCModel method. Error bars represent the standard error across participants. The translucent red region represents the task block.*

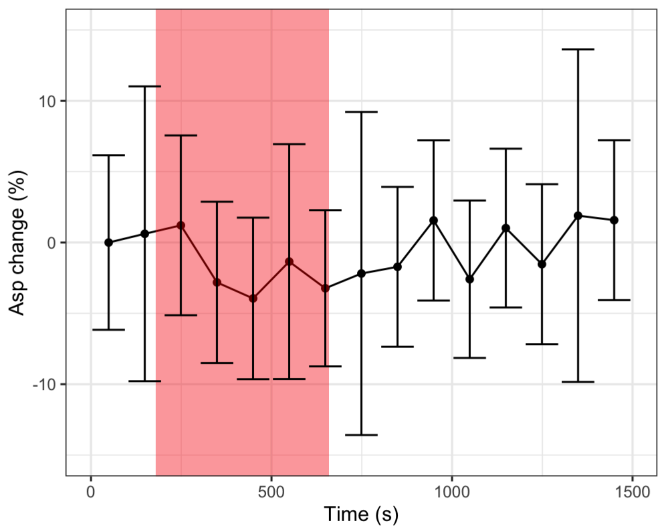


*Fig. S3 Time-courses for aspartate estimated from spectral fitting of participant averaged spectra using the ABfit method. Error bars represent the standard deviation estimated from Cramer-Rao Lower Bounds. The translucent red region represents the task block.*

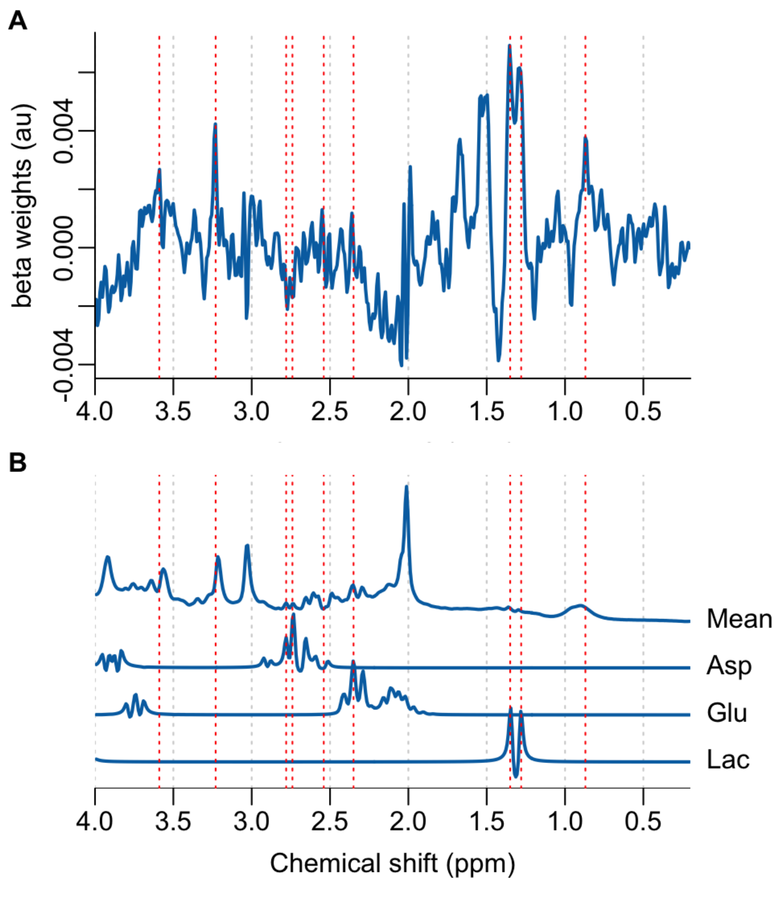


*Fig. S4 A) linear model beta weights illustrating spectral regions temporally associated with the functional task, assuming a simple boxcar function delayed by 120 seconds. B) mean fMRS spectrum with simulated glutamate (Glu), lactate (Lac) and aspartate (Asp) signals. Simulated signals are scaled to have similar maximum intensities to aid the assignment of part A). Dashed red lines highlight spectral features at 0.87, 1.28, 1.35, 2.35, 2.54, 2.74, 2.78, 3.23 and 3.59 ppm.*
